# Short-term synaptic plasticity makes neurons sensitive to the distribution of presynaptic population firing rates

**DOI:** 10.1101/707398

**Authors:** Luiz Tauffer, Arvind Kumar

## Abstract

The ability to discriminate spikes that encode a particular stimulus from spikes produced by background activity is essential for reliable information processing in the brain. We describe how synaptic short-term plasticity (STP) modulates the output of presynaptic populations as a function of the distribution of the spiking activity and find a strong relationship between STP features and sparseness of the population code, which could solve the discrimination problem. Furthermore, we show that feedforward excitation followed by inhibition (FF-EI), combined with target-dependent STP, promote substantial increase in the signal gain even for considerable deviations from the optimal conditions, granting robustness to this mechanism. A simulated neuron driven by a spiking FF-EI network is reliably modulated as predicted by a rate analysis and inherits the ability to differentiate sparse signals from dense background activity changes of the same magnitude, even at very low signal-to-noise conditions. We propose that the STP-based distribution discrimination is likely a latent function in several regions such as the cerebellum and the hippocampus.

## Introduction

The brain is a highly noisy system. At the cellular level, the neurons are unreliable in eliciting spikes and synapses are unreliable in transmitting the spikes to the postsynaptic neurons. At the network level, the connectivity and balance of excitation and inhibition gives rise to fluctuations in the background activity (Brunel, 2000; Kumar et al., 2008) which can be as high the mean stimulus response (Arieli et al., 1996). In such a noisy environment, a neuron is faced with two crucial tasks: (1) discriminating between a stimulus and fluctuations in the background activity and (2) discriminating between inputs of the same intensity (number of spikes in a period of time) but with different arrangements.

If synapses were static (that is, when the postsynaptic conductances do not depend on the immediate spike history) both these tasks cannot be accomplished, unless synapses are specifically tuned to do so. For instance, the identification of specific spiking patterns, filtering out presumed noise sequences, can be accomplished by precise tuning of synaptic weights (Gütig and Sompolinsky, 2006). This solution, however, relies on training synaptic weights using a certain supervised learning rule and even then it could only work for a specific set of spike timing sequences. Active dendrites (with voltage dependent ionic conductance) can also work as pattern detectors (Hawkins and Ahmad, 2016), but this mechanism would only work for signals constrained to locally clustered synapses. Therefore, despite being relevant for the understanding of signal processing in the brain, the mechanisms by which neural ensembles solve the activity discrimination problem have remained elusive.

Here, we show that short-term plasticity (STP) of synapses provides an effective and general mechanism to solve the two aforementioned tasks. STP refers to the observation that synaptic strength changes on spike-by-spike basis, depending on the timing of previous spikes (Stevens and Wang, 1995; Zucker and Regehr, 2002). STP arises because neurotransmitter release dynamics is history dependent. STP is believed to play several important roles in neural information processing (Fuhrmann et al., 2002; Abbott and Regehr, 2004; Rotman et al., 2011; Scott et al., 2012; Jackman and Regehr, 2017). Relevant to the two aforementioned tasks, synapses that express STP (dynamic synapses) have been suggested to help a neuron learn to identify input spike sequences (Buonomano, 2000; Rotman and Klyachko,2013). But this solution also relies on a supervised learning rule and is constrained to specific temporal sequences.

An obvious consequence of STP is that the effective postsynaptic conductances depend on the firing rates of individual presynaptic neurons. This suggests that postsynaptic targets of populations with dynamic synapses could distinguish between different input distributions even without supervised learning. To demonstrate this feature of STP, we measured the response of postsynaptic neurons for a weak stimulus with amplitude one order of magnitude smaller than the background activity. By systematically changing the distribution of rates over the presynaptic neuron ensemble, we found that weak signals can be differentiated from the noisy fluctuations if the signal is appropriately distributed over the input ensemble. The optimal distribution that maximizes the discriminability depends on the nature of STP. We found that, for facilitatory synapses, sparse codes give better discrimination between a weak signal and dense background changes of the same intensity. By contrast, for depressing synapses, sparse codes result in very negative gains in relation to dense background changes of the same intensity. We also investigated the feedforward networks with excitation and disynaptic inhibition, with the target-dependent STP, and found that this arrangement allows for extra robustness for this mechanism.

Finally, we demonstrate how STP can endow a postsynaptic neuron with the ability to differentiate sparsely encoded activity from dense activity of the same magnitude, a function that would be especially important at very low signal-to-noise regimes. Thus, our results reveal that the nature of STP may also constrain the nature of neural population code.

## Results

Here we are interested in a mechanism by which a neuronal network or a single postsynaptic neuron receiving multiple inputs may distinguish between different spike patterns with the same intensity (e.g. the same number of spikes). To illustrate this situation, consider a simple example of a sequence of 7 spikes arriving from cells presynaptic to a common readout neuron, where all the spikes can arrive from a single presynaptic neuron (Figure 1A) or from different neurons (Figure 1B). Static synapses evoke exactly the same postsynaptic conductance (PSC) sequence for the two distinct activity distributions (black lines), making both indistinguishable for a readout neuron. However, when synapses are dynamic, short-term facilitation (STF, blue line) enhances the PSC amplitudes compared to the static synapses (cf. Figure 1A,B, bottom traces). Short-term depression (STD, red line) results in a weaker response as compared to the static synapses (cf. Figure 1A,B, bottom traces). If the incoming spikes are distributed along different synapses, the sequence of PSCs is identical for all types of synaptic dynamics (cf. Figure 1A,B, bottom traces).

**Figure 1:**
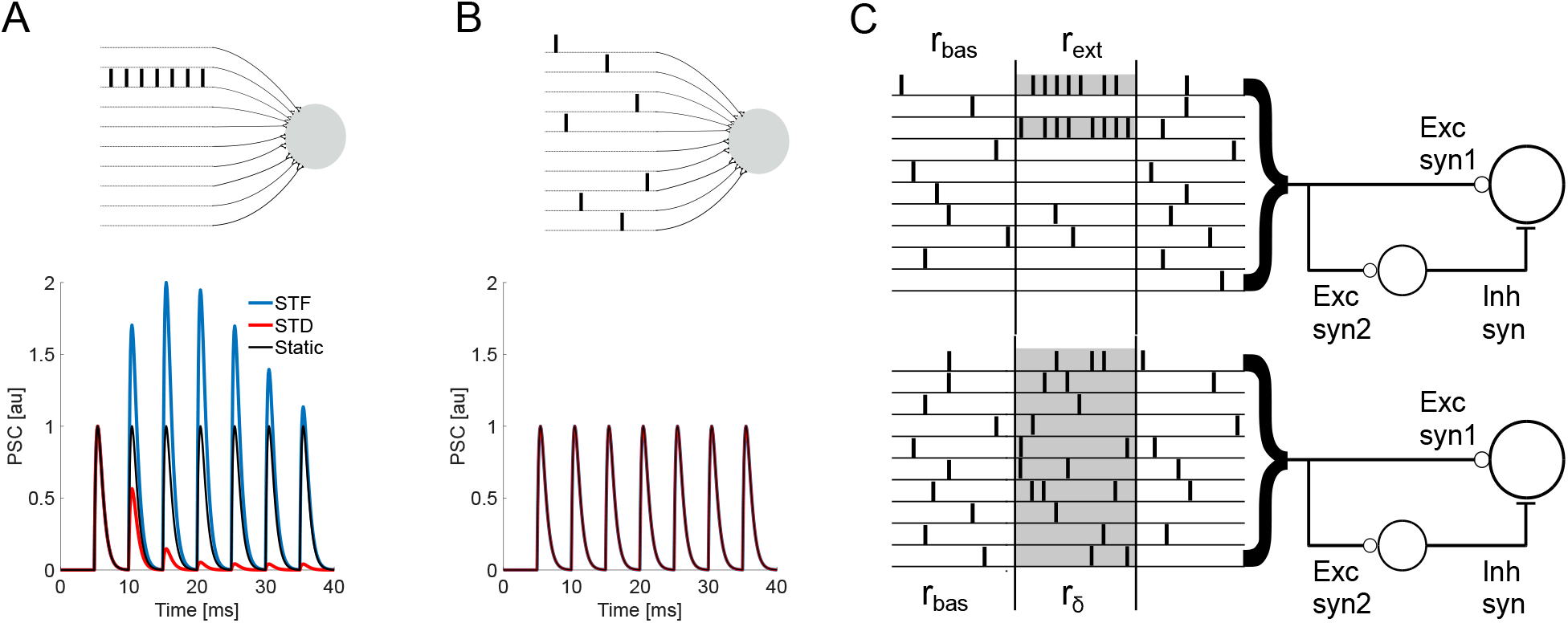
Distribution of the spiking activity over presynaptic neurons and short-term-plasticity. A Top: **A** neuron receives inputs from 10 presynaptic neurons. Of these only one of the neurons elicits 7 spikes. Bottom: The PSC generated by 7 consecutive spikes for three different types of synapses (static – black; facilitatory – blue; depressing – red). The PSCs are different for each of these three types of synapses. **B** Top: A neuron receives inputs from 10 presynaptic neurons. In this scenario each neuron elicited a single spike. But the postsynaptic neurons received 7 spikes as in the panel **A**. Bottom: The PSC generated by a sequence of 7 consecutive spikes arriving at the same time as in panel **A** coming from three different types of synapses (static – black; facilitatory – blue; depressing – red). The PSCs are identical for each of these three types of synapses (lines overlapped). C Feedforward excitation/inhibition (FF-EI) configuration and two distributions of an extra spike rate *R_ext_*. Top: the extra rate is distributed into a few presynaptic neurons (gray), with each chosen unit increasing its rate by *r_ext_* = *R_ext_/N_ext_*. Bottom: the extra rate is distributed homogeneously throughout the population of *N* units, with each unit increasing its rate by *r_δ_* = *R_ext_/N*.

In *vivo* neural coding is certainly more complex than the above example. However, this simple example suggests that in the case of a neuron receiving synaptic inputs via thousands of noisy synapses, STP could be a mechanism to differentiate between an evoked signal from the background activity fluctuations of the same amplitude, provided the former is encoded as a specific pattern that can exploit the STP properties of the synapses. In the following, we describe how well dynamic synapses could endow feedforward circuits with such activity distribution discrimination properties in low signal-to-noise regimes (Figure 1C).

### Optimal activity distribution with dynamic synapses

We implemented dynamic synapses with the rate-based TM model (Tsodyks et al., 1998) (equation 4). In this model, the instantaneous proportion of resources released (*PRR*) depend on the vesicle release probability and available vesicle resources. These variables depend on the spike history and, therefore, on the spike rate when the spikes are arriving in a Poisson manner. To quantify how the available vesicle resources change over time for a given spike rate, we defined the total amount of resources (*Q^s^*) a synapse releases over a time period of *T_s_* (equation 6; Figure 2B). Given the STP properties and a stationary baseline firing rate *r_bas_, Q^s^* reaches a stationary state 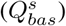. However, any transient change in the firing rate will alter the PRR (Figure 2A) and 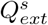.

**Figure 2:**
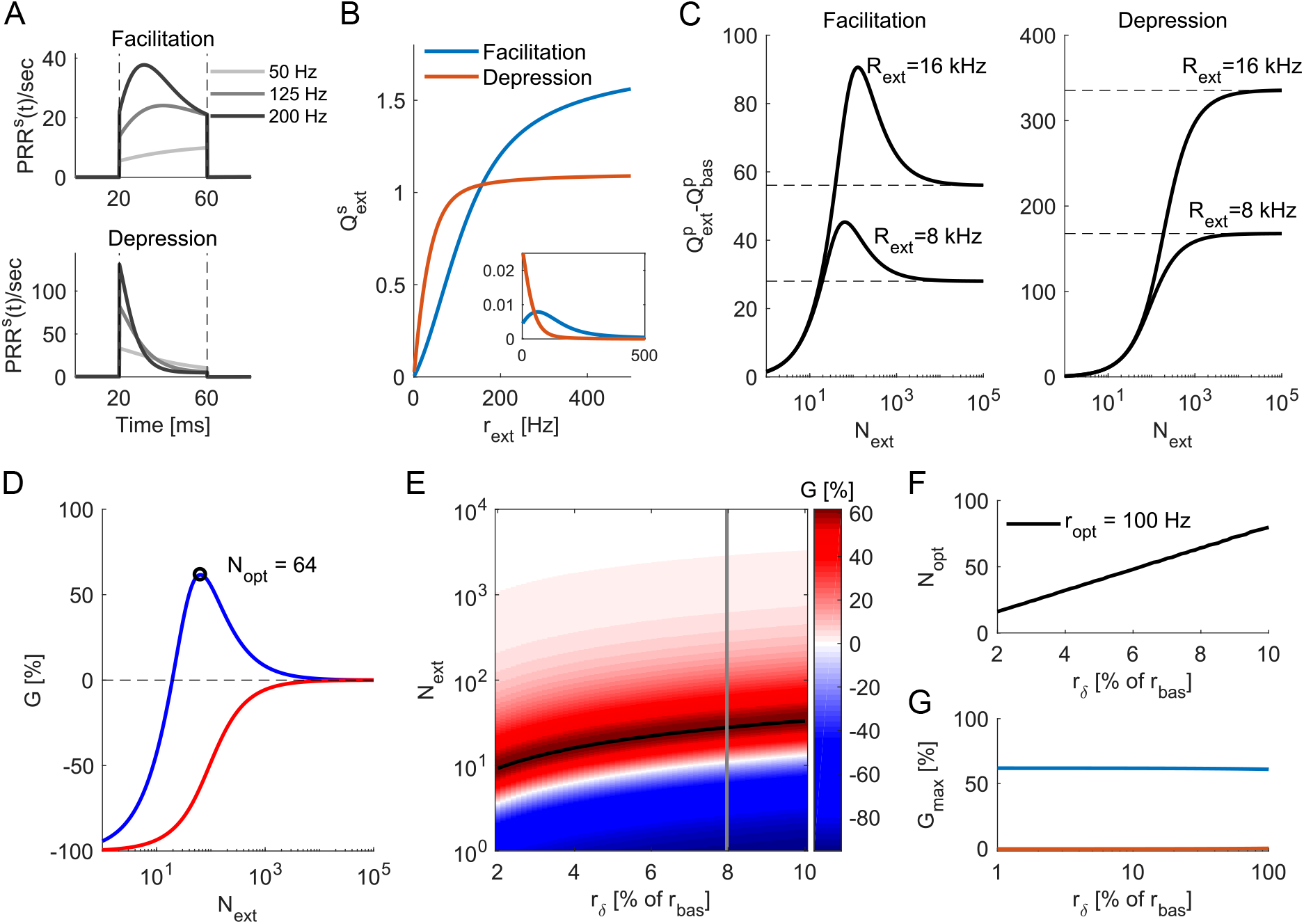
**A** Temporal profile of the *PRR* for a facilitatory (*top, U* = 0.1, *τ_f_* = 200*ms* and *τ_rec_* = 50*ms*) and depressing (*bottom, U* = 0.7, *τ_f_* = 50*ms* and *τ_rec_* = 200*ms*) synapses with increased rates during a period of *T_s_* = 40*ms*. **B** The amount of resources released by a single synapse, *Q^s^*. This was obtained by integrating *PRR^s^*(*t*) over *T_s_* (area under the curves in A, equation 6). 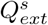 for depressing synapses saturates at lower firing rates than facilitatory synapses. The inset shows the derivative of *Q^s^* and highlights the nonlinearities in *Q^s^*, with depressing synapses showing monotonically decreasing slopes (decreasing release rate) and facilitatory synapses showing an initial region of increasing slopes (increasing release rate) with respect to *r_ext_*. **C** The extra proportion of released resources, 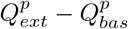 as a function of the number of presynaptic neurons (*N_ext_*) whose firing rate increases by two different values of *R_ext_*. Dashed lines mark the value achieved when *N_ext_* = *N* i.e. the dense distribution case. A population of facilitatory synapses (left) maximizes its release with low *N_ext_*, while a population of depressing synapses (right) maximizes its release with *N_ext_* = *N*. **D** The gain (G, equation 8) as a function of *N_ext_* for a fixed *R_ext_*. The *N_ext_* that maximizes *G*, for this particular extra rate, is *N_opt_* = 64 for facilitatory synapses and *N_opt_* = *N* for depressing synapses. Notice that if the extra rate is allocated in even fewer input units, *G* can be negative. **E** *G* surface for a facilitatory synapse as a function of *r_δ_* and *N_ext_*. The black line marks the maximum values of *G* i.e. *N_opt_* for each *r_δ_*. The gain curves at panel *D*, where *r_δ_* = 8% of *r_bas_*, is marked with a gray line for reference. **F** The relationship between *N_opt_* and *r_δ_* is linear. At maximal gain (*G*_max_), the firing rate of the event-related neurons (*N_ext_*) is *r_opt_* (100*Hz* for this specific example). **G** *G_max_* for the two STP regimes shown in panel *B*. For a low signal-to-basal ratio *r_δ_/r_bas_* < 1, the gain can be considered independent from the stimulus intensity *r_δ_*.

We found that the extra amount of resources released by a single synapse 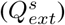 varied in a nonlinear fashion as a function of the transient rate increase *r_ext_* (equation 6, Figure 2B). The slope of 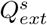 (Figure 2B *inset*) for depressing synapses is monotonically decreasing, indicating that any increase in the firing rate in those synapses will produce sublinear increase in 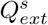, whereas for facilitatory synapses the slope initially increases, indicating that increases in the firing rate of those synapses, up to some point, will produce supralinear increase in 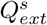.

In the brain, neurons typically receive inputs from a large ensemble of presynaptic neurons. In the ongoing activity state these neurons spike at a low-baseline firing rate (*r_bas_*) and in the event-related activity state, firing rate of a subset of presynaptic neurons is transiently increased. Therefore, it is important to understand how the total synaptic resources released (*Q^p^*) changes as the firing rate of a fraction of neurons is transiently increased. The calculation of *Q^p^* is the sum of individual *Q^s^* (equation 7). As is evident from the equation 7, *Q^p^* depends on number of neurons whose firing rate is altered. We distribute a fixed event-related population rate increase *R_ext_* into varied number of chosen synapses *N_ext_*, each of these chosen synapses increasing it’s firing rate by *r_ext_*, that is, *R_ext_* = *N_ext_ × r_ext_*, and report the changes in 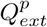.

We found that when synapses show short-term facilitation, for fixed values of *R_ext_, Q^p^* in response to external input 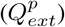 varied in a non-monotonic fashion as a function of *N_ext_* (Figure 2C left). The total synaptic resources released during the event-related activity is a sum of synaptic resources released because of the ongoing activity 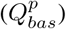 and the transient increase caused by the event 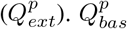 is independent of *N_ext_*, therefore, the change in synaptic resources released by a transient firing rate increase 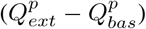 also varied in a non-monotonic fashion, as a function of *N_ext_* (Figure 2C left). By contrast, for short-term depressing synapses the change in synaptic resources released during the event-related activity varied in a monotonic fashion (Figure 2C right). For both facilitatory and depressing synapses, the change in synaptic resources released converged to their respective 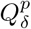 when the input rate *R_ext_* was distributed over all the input neurons such that *r_ext_* = *r_δ_* = *R_ext_/N*.

These results suggest that, when synapses are facilitatory, total synaptic resources released during a event-related activity state is maximized when event-related spiking activity is confined to a small number of synapses. 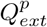 was smaller than 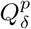 when *R_ext_* was distributed into a small subset of presynaptic neurons, because those chosen neurons spiked at very high rates and the synapses ran out of vesicle resources rapidly. When the event-related input was distributed over all the presynaptic neurons, the 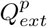 also decreased because in such a scenario *r_ext_* = *r_δ_* = *R_ext_/N* was too small to fully exploit the benefits of synaptic facilitation. In contrast to the facilitatory synapses, for depressing synapses it was more beneficial to distribute the event-related spiking activity over the whole input ensemble in order to maximize the amount total synaptic resources released. That is, in this scenario rext was small enough to avoid any losses in vesicle release because of synaptic depression.

### Activity distribution-dependent Gain

To further quantify the effect of distribution of event-related activity over the input ensemble (that is, how neurons increase their rate in the event-related phase), we defined the distribution gain *G* as the proportional change in 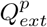 in relation to 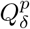 (equation 8). We found that 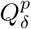 is approximately a linear function of *r_δ_* for a wide range of scenarios (see Materials and Methods) and, because of that, with the dense distribution of the inputs (when all the input neurons change their input rate by a small amount *r_δ_* in the event-related activity state), even STP type synapses behave approximately as static synapses. Therefore, *G* can be understood as a gain over a dense distribution or a gain over static synapses. For facilitatory synapses, similar to 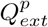, *G* also followed a non-monotonic curve as a function of *N_ext_*, with a single peak at *N_opt_* (Figure 2D, blue line). By contrast, depressing synapses resulted in negative gains to every distribution, except for *N_ext_* = *N* where *G* = 0% (Figure 2D, red line).

Next, we estimate *N_opt_* and *G* for a range of extra input intensities (Figure 2E, for facilitatory synapses). For these calculations, we parameterized the extra input *R_ext_* as a fraction of the baseline firing rate *R_bas_* (correspondingly, *r_δ_* as % of *r_bas_*, see Materials and Methods). We found that, for facilitatory synapses, *N_opt_* increased linearly with the extra input intensity (Figure 2G), resulting in an optimal encoding rate *r_opt_* which was independent of the input intensity. For depressing synapses, the optimal distribution *N_opt_* = *N* did not change with *R_ext_*, making the optimal encoding rate *r_opt_* = *r_δ_*.

Because the presynaptic neurons are assumed to be Poisson processes, an advantage of parametrize Rext in terms of fraction of *R_bas_* is that it directly translates to signal-to-noise ratio. For the example shown in Figure 2G, we found that STP could amplify the presynaptic output for weak signals (which were less than 10% of the baseline activity) by up to 60% if the extra rate was distributed over *N_opt_* synapses as opposed to N synapses. For low signal-to-noise ratios (*r_δ_* < *r_bas_*), the gain at the optimal distribution *G_max_* was approximately constant and always positive for facilitatory synapses, while depressing synapses keep *G_max_* =0 at *N_opt_* = *N* (Figure 2G). Finally, we analytically show that the independence of *r_opt_* and *G_max_* from the extra rate intensity is true for a wide range of basal rates and STP types (see Materials and Methods).

These results suggest that when synapses are facilitatory, the input should be distributed sparsely (or sparse code, that is, only a small set of neurons change their firing rate in the event-related state) to maximize the synaptic resources released at the downstream neuron. By contrast, when synapses are depressing, the input should be distributed densely (or dense code, that is, all the neurons change their firing rate in the event-related state) to maximize the synaptic resources released at the downstream neuron. Thus, for sparse population activity, facilitatory synapses get optimally utilized and depressing synapses get subutilized.

### Effects of STP parameters on optimal rate and gain

Next, we investigated how *N_opt_, r_opt_* and *G_max_* vary with STP parameters. To this end we systematically changed synapses from facilitatory to depressing by varying the set of parameters {*U, τ_f_, τ_rec_*}. We found that *r_opt_* decayed exponentially as the synapses become more depressing (Figure 3A). This follows from the fact that facilitatory synapses profit from high firing rates and depressing synapses avoid negative gains at lower rates.

**Figure 3:**
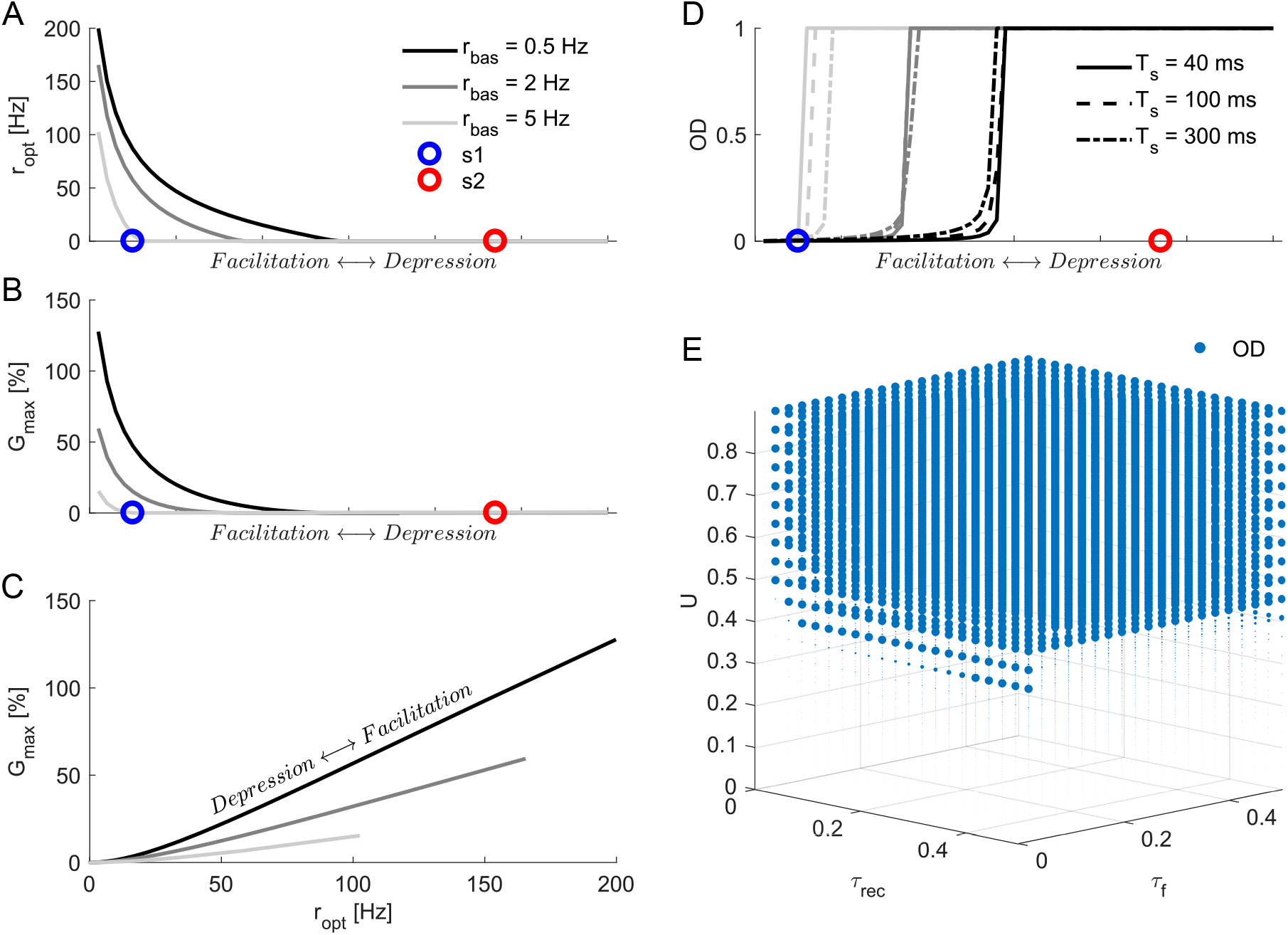
Effects of STP attributes on maximum gain of the neural population. Here the STP parameters *U*, *τ_rec_* and *τ_fac_* were varied in the following range: *U* : 0.05 → 0.9, *τ_rec_* : 20 → 500*ms* and *τ_f_* : 500 → 20*ms*. **A** The optimum frequency *r_opt_* as a function of STP properties that gradually and monotonically change the synapse from facilitatory to depressing. *r_opt_* is high for facilitatory synapses and low for depressing synapses. *r_opt_* monotonically decreases as synapses change from facilitatory to depressing. As the basal rate is increased, *r_opt_* decreased for all types of synapses. The circle markers show *r_opt_* for a facilitatory (*s*1, blue) and a depressing (*s*2, red) synapse used later in the study. **B** *G_max_* as a function of STP properties that gradually and monotonically change the synapse from facilitatory to depressing. **C** The relationship between *G_max_* and *r_opt_*. Notice the approximately linear relationship for facilitatory synapses, with the slope steadily decreasing with increasing *r_bas_*. **D** Optimal distribution of rate over *N_opt_* presynaptic neurons that maximize the gain *G*. The change from sparse to dense optimal distribution (0 to 1 *OD*, respectively) is abrupt and occurs approximately at the same STP region for all *T_s_*. However, the transition point where the input distribution changes from sparse to dense code is strongly modulated by *r_bas_* – higher basal rates allows for sparse code only for more facilitatory synapses. **E** Optimal distribution of rate as a function of the three key model parameters (*U*, *τ_rec_* and *τ_fac_*). The variable U is the most influential in defining the optimal encoding distribution, with *U* ~ 0.45 defining the OD transition point for *T_s_* = 40*ms* and *r_bas_* = 0.5*Hz*. Marker sizes represent *OD* values, with large ones for *OD* = 1 and small ones for *OD* ≈ 0.

The maximum gain *G_max_* also decreased exponentially as synapses were systematically changed from facilitatory to depressing (3B). We found that the relationship between gain and optimal rate was linear from mildly to strongly facilitatory synapses (3C), with larger basal rates constraining the optimal conditions to lower rates with lower gains.

Interestingly, increasing the basal firing rate *r_bas_* substantially reduced *r_opt_* and *G_max_*. This is surprising because, at such low values of spiking rates, STP effects are hardly perceivable in traditional paired-pulse ratio analyses. The high value of *G_max_*, when the system operates at low *r_bas_*, is because of the many synapses taking advantage of the nonlinearities in their individual *Q^s^*. That is, even if there was a small change in *Q^s^*, because of the large number of input synapses, the collective effect was much higher. Therefore, increased baseline activity strongly impaired the capacity to exploit these nonlinearities and, ultimately, the distinction of activity distributions (sparse vs. dense).

### Relationship between facilitatory synapses and sparse coding

We quantified the optimal distribution of an evoked neural signal by *OD* (see Materials and Methods). High *OD* (*OD* → 1) indicates a dense distribution in which many neurons spike to encode the extra activity, whereas low *OD* (*OD* → 0) implies a sparse distribution. We found that *OD* changed abruptly from sparse to dense as synapses were changed from facilitatory to depressing (Figure 3D). Facilitatory synapses yielded maximum response for sparse while depressing synapses yielded maximum response (avoid negative gains) for dense distributions. The transition point from sparse to dense OD did not depend on the stimulus duration. However, the basal rate strongly modified the transition point, with higher *r_bas_* allowing only strongly facilitatory synapses to take advantage of sparse distributions. This configuration remained independent of the stimulus intensity as long as the circuit operates at low signal-to-noise conditions (*r_δ_/r_bas_* < 1, Figure 2G).

In the above, we characterized the synapses as facilitatory and depressing without being specific about the model parameters. Next, we systematically varied each of the STP parameters and measured the OD for maximum gain. We found that the transition region was primarily governed by the facilitation factor *U* (*U* ~ 0.45), with a weak dependence on *τ_rec_* and *τ_f_* (Figure 3E). The relative contribution of *τ_rec_* and *τ_f_* became more relevant at higher *T_s_* (Supplementary Figure 1).

These results clearly highlight the importance of the stationary basal rate in how well the synaptic gain modulation operates, as only low *r_bas_* allows for significant gains. Importantly, the switch-like behavior of the optimal distribution indicates that, for a given population code, there is a robust range of STP attributes that could produce positive gains. This transition point seems to be relatively independent of the signal duration but is strongly affected by *r_bas_*. Finally, having a low initial release probability (defined in the model by a low *U*) seems to be the preeminent feature in defining the optimal *OD*. Next, we investigate how fine tuning of parameters could affect the optimal conditions.

### Effects of different sources of enhancement on *G_max_*

The enhancement of the output at facilitatory synapses could, in principle, have many causes (Valera et al., 2012;Thanawala and Regehr, 2013; Jackman and Regehr, 2017). Using the TM model (equation 4), we phenomenologically accounted for two important sources: a low initial release probability which sequentially increases with each incoming spike (Jackman et al., 2016) and fast replenishment of readily available resources (Crowley et al., 2007). The first characteristic is mimicked by a low facilitation factor *U*, which determines the initial release probability after a long quiescent period and the proportional increase in it after each spike. The second mechanism is captured by a fast recovery time constant *τ_rec_*.

We systematically varied *U* and *τ_rec_* and measured *G_max_* and *r_opt_*. We found that several different combinations of *U* and τ_rec_ resulted in the same optimal distribution gain and rate. However, when we changed *U* and *τ_rec_* while keeping the *r_opt_* fixed, *G_max_* could no longer be kept constant and vice versa. For instance, the two parameter sets {*U* = 0.05, *τ_rec_* = 90*ms*} and {*U* = 0.1, *τ_rec_* = 15*ms*} gave *r_opt_* = 150*Hz* (Figure 4A), but the first parameter set gave *G_max_* = 109% and the second parameter set gave *G_max_* = 92% (Figure 4B). Holding U fixed and choosing *τ_rec_* to match with different *r_opt_* showed that *G_max_* consistently dropped for higher *U* (Figure 4C,D).

**Figure 4:**
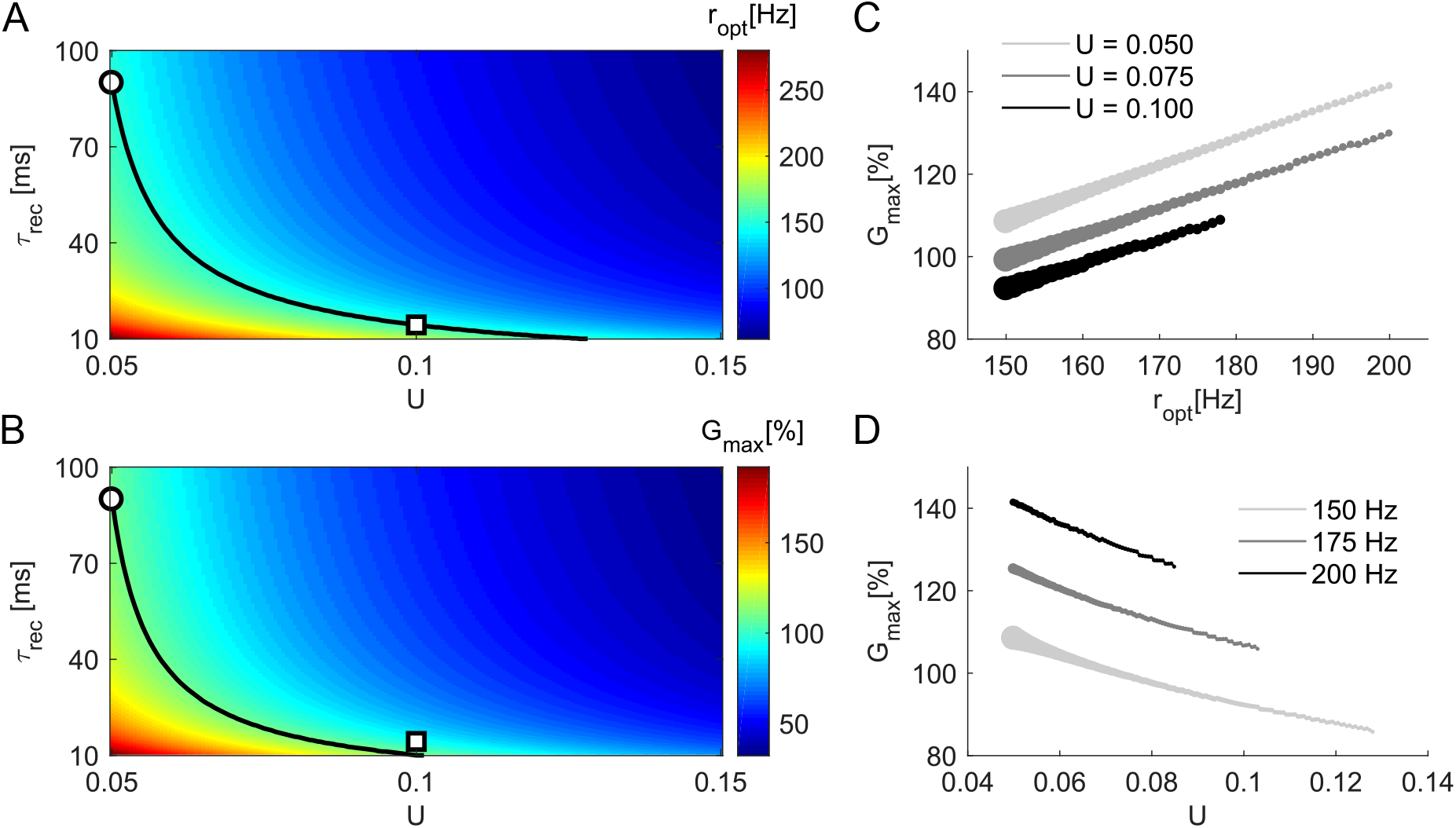
Effects of resources recovery time *τ_rec_* and facilitation factor *U* on *G_max_* and *r_opt_* for *T_s_* = 40*ms*, for facilitatory synapses. **A** *r_opt_* as a function of facilitation factor (*U*) and recovery time constant (*τ_rec_*). The *r_opt_* surface shows that a given optimum encoding rate can be matched by different combinations of synaptic parameters. For example, the iso-frequency curve of 150*Hz* (black line) is achieved with *U* = 0.05 and *tau_rec_* = 90*ms* (∘) or with *U* = 0.1 and *τ_rec_* = 15*ms* (□). **B** *G_max_* as a function of facilitation factor (*U*) and recovery time constant (*τ_rec_*). Same maximal gain can observed for many different combinations of *U* and *τ_rec_*. The black line shows the contour for *G_max_* = 109%. The two configurations with same *r_opt_* marked in panel A have distinct gains (∘ = 109%, □ = 92%). **C** We fix *U* and vary *τ_rec_* (circle sizes) to match *r_opt_* (x axis), then observe the gain. Larger values of *U* systematically produce smaller gains. Recovery time has a lower boundary *τ_rec_* = 10*ms*. **D** *G_max_* as a function of *U* for three different values of *r_opt_*. Larger values of *U* require smaller values of *τ_rec_* (circle sizes) to match the same *r_opt_*, but as a consequence the gain decreases as we increase *U*.

These results indicate that, in terms of maximum gain *G_max_*, the fine tuning of intracellular mechanisms that work to steadily increase a low initial release probability might be more important than fast vesicle replenishment mechanisms. This remains true for larger Ts (Supplementary Figure 2).

In summary, these results show that a set of presynaptic STP parameters generates a gain surface *G* that, in principle, could be tuned to match presynaptic population activity characteristics. The optimum rate and the maximum gain are independent of the stimulus intensity for a low signal-to-noise ratio, with facilitatory synapses yielding high gains for sparse distributions while depressing synapses avoid negative gains only with dense distributions. For low basal activity (*r_bas_* = 0.5*Hz*) and short duration integration window (*T_s_* = 40*ms*) conditions, the parameter *U* is the principal determinant of the optimal distribution. Furthermore, lower *U* yields a higher gains than lower *τ_rec_* when the optimal encoding rate is kept constant.

### Feedforward inhibition and heterogeneous STP

In the above we ignored the fact that presynaptic STP can be target-dependent (Reyes et al., 1998; Markram et al., 1998;Rozov et al., 2001; Sun et al., 2005; Pelkey and McBain, 2007; Bao et al., 2010; Blackman et al., 2013;Larsen and Sjöström, 2015;Éltes et al., 2017) and the spike trains coming from the same axon can be modulated by different short-term dynamics at different synapses. In the following, we describe the effects of such heterogeneity in a feedforward excitation and inhibition (FF-EI) motif (Figure 1C), an ubiquitous circuit motif across the brain (Klyachko and Stevens,2006; Dean et al., 2009; Isaacson and Scanziani, 2011; Wilson et al., 2012; Jiang et al., 2015; Grangeray-Vilmint et al.,2018).

We extend our previous analysis to a scenario in which the presynaptic population makes synaptic contacts not only with a readout neuron, but also with the local inhibitory population which projects to the readout neuron creating the FF-EI motif. Both, the readout neuron and the inhibitory group receive the same spike trains via two different types of synapses, *s*1 and *s*2 (Figure 1C). Because the presynaptic population activity is the same for both synapses (*r_bas_* = 0.5*Hz*, *T_s_* = 40*ms*), the differences in gain (*G*) are governed by the STP properties of the two synapses. Figure 5A shows *G* for a facilitatory (*s*1, *U* = 0.1, *τ_f_* = 200*ms*, *τ_rec_* = 50*ms*) and a depressing (*s*2, *U* = 0.7, *τ_f_* = 50*ms*, *τ_rec_* = 200*ms*) synapse.

**Figure 5:**
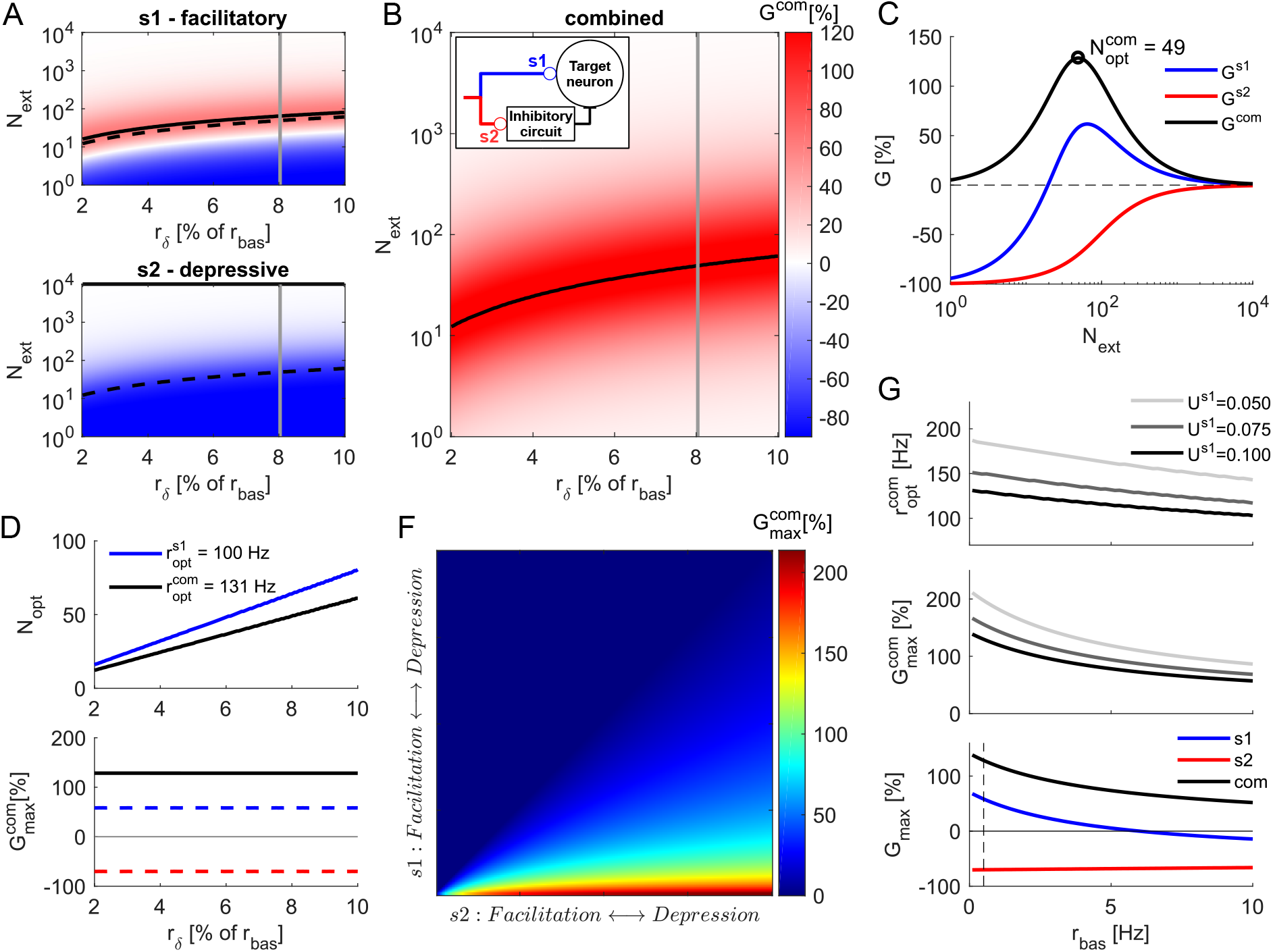
Combined optimal distribution of input in a feedforward excitation-inhibition circuit with target-dependent STP. **A** *G* as a function of *r_δ_* and *N_ext_* for a facilitatory synapse (*s*1, *top*) and for a depressing synapse (*s*2, *bottom*). This is similar to the Figure 2E. **B** The combined gain (*G^com^* = *G*^*s*1^ − *G*^*s*2^) of the FF-EI circuit as a function of *r_δ_* and *N_ext_* obtained by combining the gains of the feedforward excitation and inhibition branches. The black line marks the 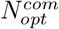 for every stimulus intensity *r_δ_* and is represented with dashed black lines in panel *A*. In-box: schematic of the FF-EI circuit. C Gain as a function of *N_ext_* for *r_δ_* = 8% of *r_bas_* (gray lines in panels *A* and *B*). *G^com^* inherits the non-monotonicity from *G*^*s*1^(*blue*, compare with Figure 2D). The gain for a depressing synapse is negative (*G*^*s*2^, *red*) for every *N_ext_* < *N*. **D** *top N_opt_* as a function of *r_δ_* produces iso-frequency lines (compare with Figure 2F). 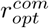 is markedly larger than 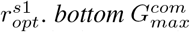 is independent of *r_δ_* (for *r_δ_* < *r_bas_*, compare with Figure 2G). Gain for both synapse types at 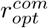 (dashed lines). The small decrease in synaptic gain for *s*1 is compensated by putting *s*2 in a very negative gain region. **E** 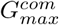 surface for different combinations of STP characteristics of *s*1 and *s*2. Notice that 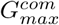 steadily increases for *s*1 → Fac or *s*2 → *Dep*. **F** Effects of ongoing basal activity *r_bas_* on optimal conditions for *T_s_* = 40*ms*. Increasing basal activity decreases the combined optimum rate. Results for 3 different *U* at the facilitatory synapse (*top*). Increasing basal activity consistently decreases *G_max_* (*bottom*). 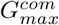 decay happens mostly due to decay of the positive gain at the facilitatory synapse *s*1 (*blue*), while the negative gain at the depressing synapse *s*2 is kept negative and change only slightly (*red*). Dashed vertical line marks the basal activity used for most part of our analysis, *r_bas_* = 0.5*Hz*, where both branches contribute significantly to increase the combined gain.

In the case of a FF-EI network, those two synapse types may be associated with the two branches, for example *s*1 to the feedforward excitation (FFE) branch (targeting a principal neuron) and *s*2 to the feedforward inhibition (FFI) branch (targeting local interneurons which eventually project to principal neurons) (Figure 5B inset). In this arrangement, the combined gain is determined by the two branches *G^com^* = *G*^*s*1^ − *G*^*s*2^. We found that the combined gain of the FF-EI circuit also varied non-monotonically as a function of *N_ext_* and peaked at 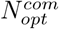 which corresponded to the combined optimum encoding rate 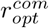 (Figure 5B,C). Note that the combined maximum gain of the FF-EI circuit is larger than the gain obtained via the FFE branch with facilitatory synapses alone (Figure 2C). This substantial increase is a consequence of the strictly negative profile of *G*^*s*2^. When the extra input is distributed in 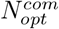 units (sparse coding), the depressing branch of the FF-EI drove the local inhibitory group with weaker strength than a scenario in which *N_ext_* = *N* (dense coding). Therefore, with sparse distribution of the input the readout neuron experienced strong excitation from the FFE branch and weak inhibition from FFI branch.

Similar to the behavior of facilitatory synapses, in the FF-EI network 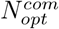 increased linearly as a function of *r_δ_*, maintaining a constant optimal encoding rate 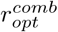 (Figure 2D, top). We also observed that 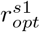 was larger than 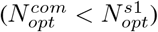, making the isolated gain of *s*1 suboptimal. However, this can be compensated by putting *s*2 into a very negative gain region (Figure 5D *bottom*, red dashed line), with a sparse distribution of the inputs. We show analytically that 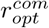 and 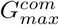 are independent of the extra rate for a wide range of conditions (see Materials and Methods).

We extended this analysis to a large range of {*s*1, *s*2} STP combinations by gradually changing the set of parameters {*U*, *τ_f_*, *τ_rec_*} (Figure 5F). We found that 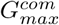 increased monotonically when we made the synapse *s*1 more facilitatory or when we made the synapse *s*2 more depressing. The anti-diagonal (where *s*1 = *s*2) marked the region of zero gain and any point above it (*s*2 more facilitatory than *s*1) resulted in 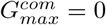, whereas any point below it (*s*1 more facilitatory than *s*2) resulted in 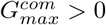. As expected, sparse distribution of the extra rate resulted in very high gain when *s*1 is highly facilitatory and *s*2 highly depressing.

### Effects of basal activity on the FF-EI network

Next, we investigated the effects of the stationary basal activity at the combined optimal conditions of a FF-EI network. We found that the optimal rate and optimal gain both decreased as *r_bas_* was increased (Figure 5G). Separation of the individual contributions of *s*1 and *s*2 branches revealed that this decrease was primarily because of a reduction in the gain of facilitatory synapses (*s*1) whereas the strong negative gain of depressing synapses (*s*2) remained approximately unaltered. This suggests that a population of facilitatory synapses will rapidly lose its activity distribution discrimination capacity as the baseline firing rate is increased, whereas a population of depressing synapses can preserve this capability even at larger basal rates.

Thus, these results show that a FF-EI network with target-dependent STP can make the discrimination of sparse inputs more robust than what could be achieved by the feedforward excitation alone. This can be achieved when the excitatory branch is facilitatory while the activation of the inhibitory branch is depressing (by placing *s*1 and *s*2 at the region below the anti-diagonal on Figure 5F).

### Sparse code identification by a postsynaptic neuron model

The ability of STP to amplify the output of a presynaptic population would be functionally relevant only if this amplification is transferred to the postsynaptic side. We tested the postsynaptic effects of the STP based modulation of the presynaptic activity distribution by simulating an integrate and fire (IF) neuron model (equation 17) as a readout device for a FF-EI circuit (Figure 6A). We simulate a presynaptic population with characteristics similar to the cerebellar molecular layer, a massively feedforward system with properties much alike the ones we have described so far (Ito,2006).

**Figure 6:**
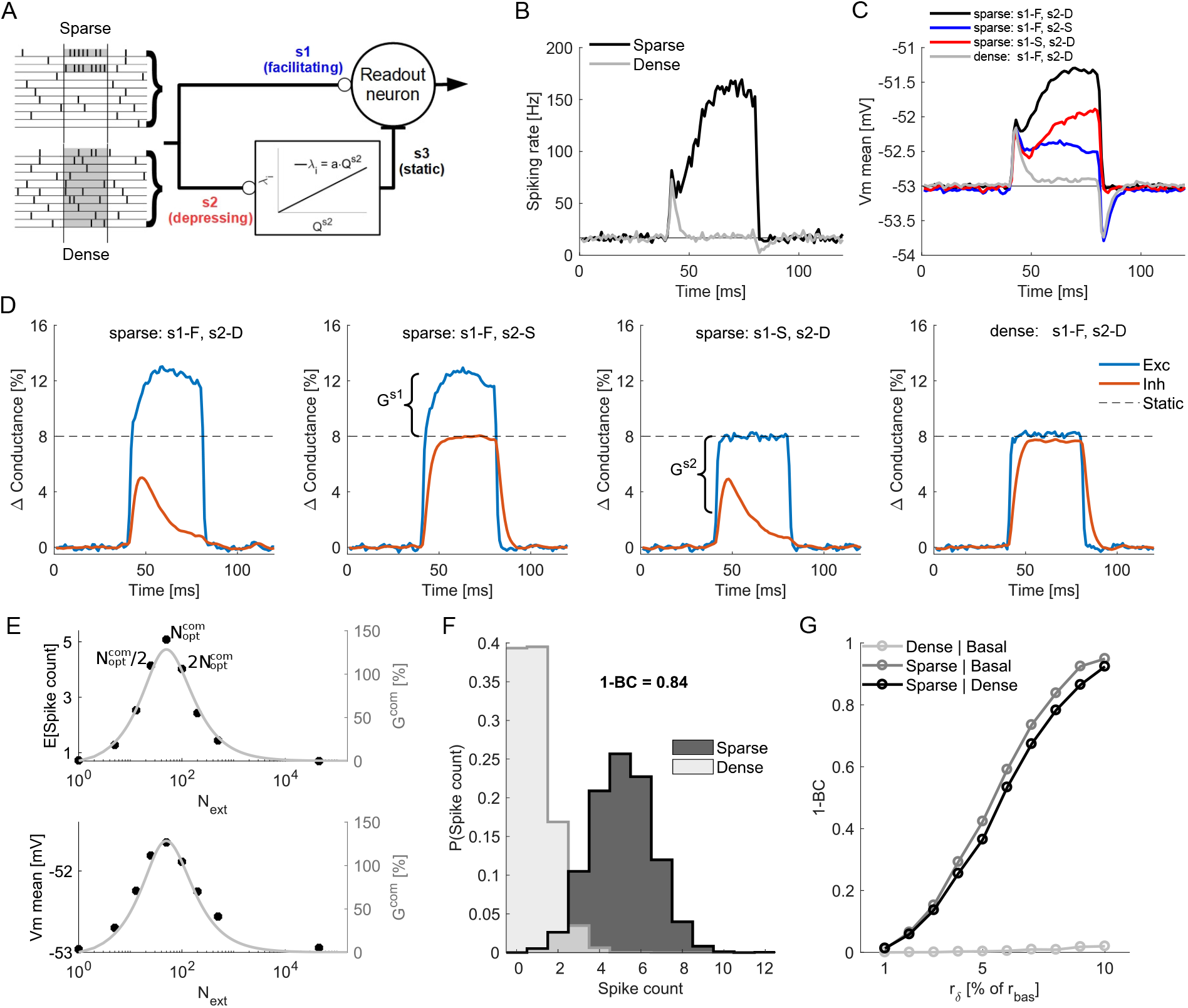
Transfer of STP gain from presynaptic side to postsynaptic neuron. **(A) Right** Schematic representation of a readout neuron receiving feedfoward excitation from *N* = 160, 000 neurons and feedforward inhibitory input. The inhibitory group was driven by the PRR of synapses of type *s*_2_ according the the linear function shown in the rectangular box. **(A) Left** Sparse and dense input patterns are schematically shown. The basal rate was set to *r_bas_*= 0.5*Hz*. The population temporally (*T_s_* = 40*ms* marked in gray) increases its firing rate by *R_ext_* = 1…10% of *R_bas_* in two different configurations: sparse (Top, *N_ext_* = *N_opt_*) or dense distribution (Bottom, *N_ext_* = 16,000). For panels *B, C, D* and *E, R_ext_* = 8% of *R_bas_*. **B)** PSTH of the spiking rate (3000 realizations) of readout neuron receiving sparse (black) or dense (gray) distribution of input activity. **C)** Mean membrane potential for sparse input (*black*) and dense input (*gray*) when the synapse si was facilitatory and synapse *s*_2_ was depressing. Red trace: Membrane potential when the synapse *s*_1_ was facilitatory and synapses *s*_2_ was static. Blue trace: Membrane potential when the synapse *s*_1_ was static and synapses *s*_2_ was depressing. The red and blue traces show the contributions from synaptic facilitation (on the feedforward excitation branch) and depression (on the feedforward inhibitory branch) to the neuron response. **D)** Changes in the total excitatory and inhibitory conductances for the four configurations of synapses (as show in the panel **C.** The dashed line marks the conductance changes for the static synapses condition. **E)** Effect of varying *N_ext_* as a proportion of 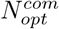 on the expected spike count (*top*, black circles) and the mean membrane potential (*bottom*, black circles) during the event-related activity period. Both profiles match the combined gain curve (gray line, compare with Figure 5C), with peak at 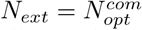. **F)** Probability distribution of output spike counts within T_s_. The sparse distribution increases substantially the elicited number of spikes at the readout neuron when compared to the dense distribution. **G)** Separation (1-BC) between spike count distributions as a function of *r_δ_*. The sparse distribution produced increasingly substantial separation when compared to basal (*dark gray*) and dense distribution (*black*) whereas the separation was always small when comparing dense distribution with basal activity (*light gray*).

Specifically, the readout neuron received input from 160,000 presynaptic neurons. The presynaptic background activity was modeled as independent and homogeneous Poisson spike trains with average firing of *r_bas_* = 0.5*Hz* (*R_bas_* = 80*kHz*). In addition, the population of presynaptic neurons increased their firing rate (*R_ext_* = 1 … 10% of *R_bas_*) during a brief time window (*T_s_* = 40*ms*) to mimic a event-related activity. The extra presynaptic activity was either confined to a small set of presynaptic neurons (*N_ext_* = *N_opt_*, *sparse*) or distributed over a large number of neurons (*N_ext_* = 16,000, *dense*). The excitatory synapses onto the readout neuron (*s*1) were facilitatory and the STP parameters for each synapse were drawn from a Gaussian distribution (*s*1, *U* : *mean* = .1, *s.d*. = .02, *τ_rec_* : *mean* = 50, *s.d*. = 10*ms* and *τ_f_* : *mean* = 200, *s.d*. = 40*ms*). The feedforward inhibitory activity was modeled as a Poisson process whose firing rate (*λ_i_*, equation 19) was linearly dependent on the excitatory input of depressing synapses, whose STP parameters for each synapse were drawn from a Gaussian distribution (*s*2, *U* : *mean* = 0.7, *s.d*. = .14, *τ_rec_* : *mean* = 200, *s.d*. = 40*ms*, and *τ_f_* : *mean* = 50, *s.d*. = 10*ms*).

The distribution of the input had a noticeable effect in the output of the target neuron, as shown by the peristimulus time histogram (Figure 6B). While the dense distribution elicited transients at the beginning and ending of the stimulus period because of the inhibition slow time constant, the sparse code elicited a sustained elevated firing rate response throughout the stimulus period. The stimulus induced membrane potential responses for the two types of input patterns (dense and sparse) were also similar to the firing rate responses (Figure 6C). By interchangeably setting *s*1 and *s*2 to static, we identified that both branches contributed significantly to keep the mean membrane potential high in the presence of extra sparse input.

The contribution of each branch becomes clear at the average change in the total excitatory and inhibitory conductances of the readout neuron. When both synapses were dynamic and the stimulus was sparse (Figure 6D, *leftmost*), the average excitation was larger (because of synaptic facilitation) and the average inhibition was lower (because of synaptic depression) than the average changes caused by a stimulus of the same intensity but with dense distribution (Figure 6D, *rightmost*). Note how, with dynamic synapses and dense distribution of the stimulus, the conductance changes matched the expected change for static synapses (dashed line). When we kept the stimulus distribution sparse, but interchangeably set *s*1 and *s*2 to static, the conductance trace related to the static branch reached the same value as for the dense distribution and the system was left with the gain produced at the dynamic branch. Dense distributions, therefore, do not exploit the STP nonlinearities and the synapses behave approximately as static, as predicted.

Next, we systematically changed *N_ext_* as percentages of *N_opt_* (*N_ext_* = 1,10,25,50,100, 200,400,1000% of *N_opt_*, black circles in Figure 6E) and found that both the mean membrane potential and the average spike count during the stimulus period followed profiles that closely matched the predicted *G^com^* curve (Figure 6E). This result confirms that the modulation of the proportion of released resources from the presynaptic population is faithfully translated into postsynaptic variables (gain estimated as the presynaptic population and membrane potential and spike rate measured on the postsynaptic neuron side). Furthermore, this result also highlights the robustness of this mechanism – even with considerable deviations from the optimum encoding distribution (*N_ext_* = 50% or *N_ext_*= 200% of *N_opt_*, marked as the first black points at left and right from *N_ext_* = *N_opt_*), the evoked responses remained reasonably close to the optimal.

To further assess how individual realizations of the sparse input could be distinguished from a dense input of the same intensity, we sampled the output spike count of the readout neuron for a period of 40*ms* during the ongoing basal activity just before the stimulus and during the 40*ms* stimulus period for both sparse and dense distributions (Figure 6F). We used the Bhattacharyya coefficient (BC) as a measure of overlap between these sample distributions and 1 – *BC* as a measure of difference (Figure 6G). The dense input had almost complete overlap with the basal condition. On the other hand, the sparse input produced increasingly different response distributions from both the dense input and basal condition, with almost complete separation at *r_δ_* = 10% of *r_bas_*.

Taken together, these results illustrate the potential role of dynamic synapses in amplification of sparse signals at the presynaptic side (*Q^p^, G*), even when such signal intensity is just a small fraction of the ongoing basal activity and, therefore, likely to be buried in proportionally large noise fluctuations. In addition, for a dense distribution of the input, the system can preserve short periods (~ 10*ms*) of increased (decreased) spike probability right after stimulus onset (offset) due to delayed inhibition, which is a known characteristic of FF-EI motifs and might serve as indication of global background rate changes.

### Continuous rate distribution

Thus far, we have considered a binary distribution of the extra rate: a fraction of presynaptic cells increased their rate by *r_ext_* or not at all. Although some bursting networks (e.g. cerebellar parallel fibers) do seem to operate in a quasi-binary fashion (burst or no-burst), it is important to extend the analysis to continuous distributions, which most parts of the brain seem to operate under. We do this by assuming that the distribution of event-related neural firing rates follows a Gamma distribution, which allows us parameterized control of the sparseness of the neural code (with the mean of the distribution) and of the distribution shape (with the skewness and kurtosis):

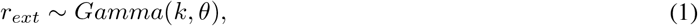

where *k* is the shape parameter and *θ* is the scale parameter. For *k* = 1 it is equivalent to an exponential distribution and, for increasing values of *k*, it becomes a right-skewed distribution, with the skewness approaching zero for higher values of *k* (becoming approximately Gaussian). For each shape parameter, we controlled the mean of the distribution by varying the scale parameter, because for a Gamma distributed *r_ext_* the expected value is

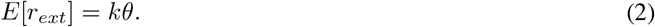

For the above specified distribution of extra rates and a given presynaptic set of STP parameters, the expected amount of resources released by a population is

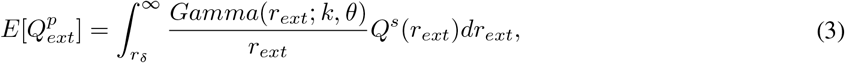

which we solved numerically for two synapse types (*s*1-facilitatory and *s*2-depressing) and a range of rate distributions (Figure 7B). The distribution gain *G* for 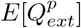 was then calculated in relation to the dense case, where *N_ext_* = *N* and *r_ext_* = *r_δ_* (Figure 7A).

**Figure 7:**
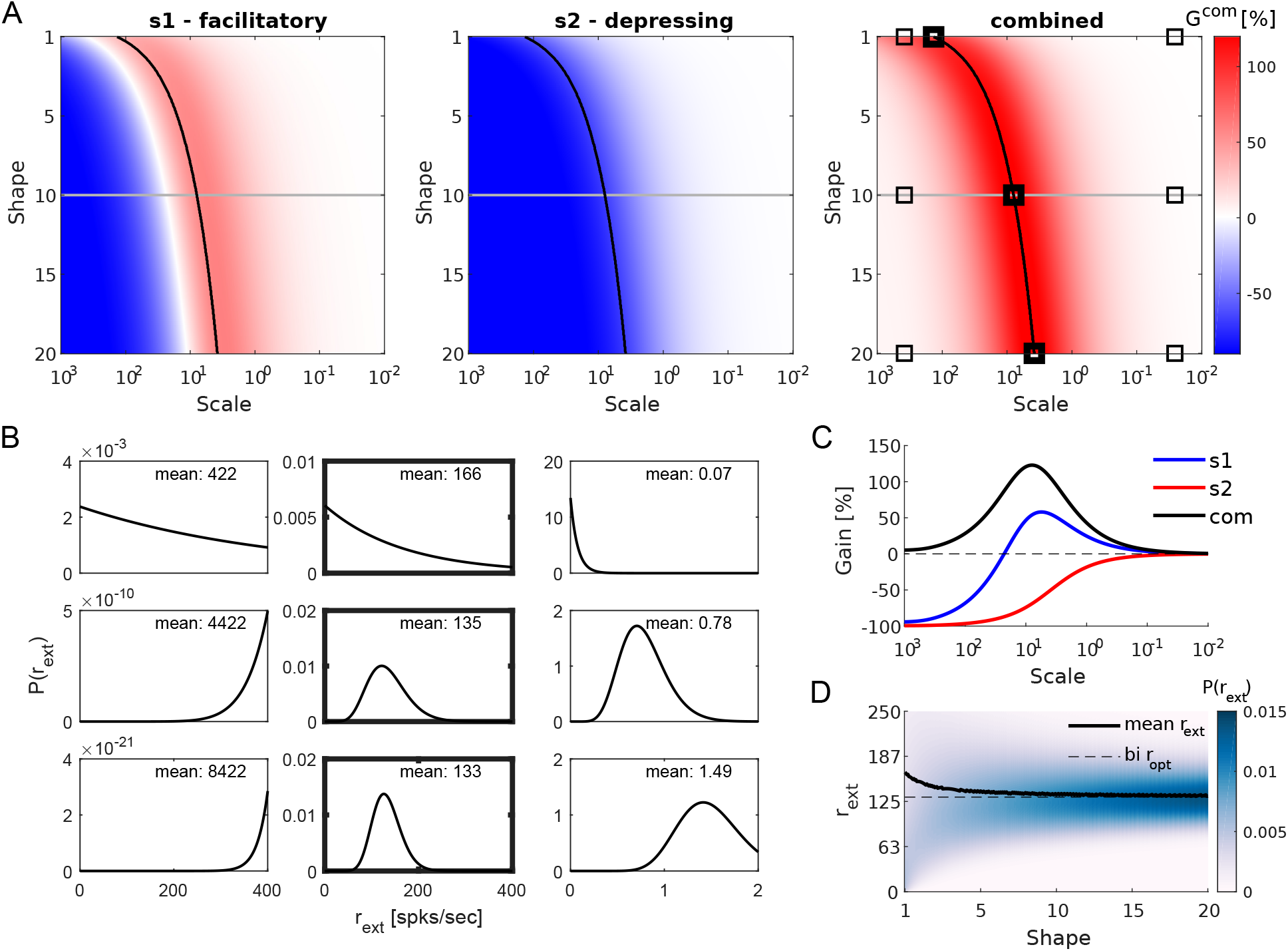
Synaptic gain when firing rates of individual neurons (*r_ext_*) were draw from continuous distributions (Gamma distribution). **A** Gain surfaces for a facilitatory synapse (*left*), for a depressing synapse (*middle*) and for the combined effect in a FF-EI (*right*). Increasing the shape parameter moved the distribution from an exponential to a right-skewed to an approximately Gaussian one. Decreasing the scale parameter moved the distribution from a high mean and high variance (sparse) to a low mean and low variance one (dense). Black lines mark the *θ* that resulted in maximum combined gain for each value of *k*. Similar to the binary distribution, the 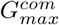 is obtained by putting *s*1 in positive and *s*2 in negative gain regions. Note that the gain values are in the same range as in Figure 5A,B. **B** Representative examples of Gamma functions used to model the distribution *r_ext_*. The nine examples correspond to the points marked in panel **A**. Changing *k* affects the shape of the distribution: exponential (*top row*), right-skewed (*middle row*) and approximately Gaussian (*bottom row*) shapes. Changing *θ* affects the scale of the distribution (sparsity of the population code): high mean and variance (*left column*), optimal mean and variance (*middle column*) and low mean and variance (*right column*). **C** Gain curves for a facilitatory synapse (*blue*), for a depressing synapse (*red*) and for the combined effect in a FF-EI (*black*). These curves were obtained for a fixed *k* = 10 (gray lines on panel **A**) and gradually changing *θ*. The gains as a function of the Gamma activity distribution follow a profile similar to the binary distribution (compare with Figure 5C). **D** Gamma-distributed *r_ext_* (color plot) as a function of shape parameter. Mean *r_ext_* from the Gamma distributions obtained with the optimal *θ* for each value of *k* (black lines on panel A). As the Gamma shape moves from an exponential to a Gaussian one (increasing *k*), the mean of the optimal distribution approaches the *r_opt_* for the binary distribution.

We found that, similar to the binary distribution case, the gain for facilitatory synapses followed a non-monotonic curve as a function of *θ* (for a fixed *k*), with negative values at high *θ* (overly sparse distribution), a single peak at the optimal *θ* choice and convergence to 0 at low *θ* (dense distribution). By contrast, depressing synapses showed negative gains, monotonically converging to zero at low *θ*. The combined gain reached high values when *s*1 synapses were in very positive and *s*2 synapses were in very negative operating regions (Figure 7C).

Interestingly, not only the gain magnitudes were very similar to the ones obtained with binary distributions (compare colorbars of Figure 5A,B and Figure 7A), but also with continuously distributed rates the points of maximum gain were obtained at high mean rates (in relation to *r_δ_*) and, therefore, representative of sparse distributions of the population activity. For increasing values of *k*, the skewness of these distributions approached zero (i.e. became closer to a Gaussian) and the mean *r_ext_* of the optimal *θ* approaches the *r_opt_* obtained by binary distributions. These results further highlight the relevance of the activity distribution-dependent gain modulation in presynaptic populations with STP.

## Discussion

Our results suggest a close relationship between short-term synaptic plasticity and the nature of the population code that, within physiological values, can endow a postsynaptic neuron with the ability to discriminate between weak signals and background activity fluctuations of the same amplitude.

### Relevance to specific brain circuits

We have shown that STP can enhance the effective input when (1) stimulus is sparse, temporally bursty and (2) feedforward excitatory synapses on the principal cells are facilitatory and feedforward excitatory synapses on local fast-spiking, inhibitory interneurons are depressing. These two conditions are fulfilled in several brain regions.

In the cerebellum, glomeruli in the granular layer actively sparsify the multimodal input from mossy fibers into relatively few simultaneously bursting parallel fibers (PF) (Billings et al., 2014) projecting to Purkinje cells (PuC). A single PuC might sample from hundreds of thousands of PFs (Tyrrell and Willshaw, 1992; Ito, 2006). In behaving animals, PF present two stereotypical activity patterns, a noisy basal state with rates lower than 1*Hz* during long periods interleaved by short duration (~ 40*ms*), high frequency (usually > 100*Hz*) bursts carrying sensory-motor information (Chadderton et al., 2004; Jorntell and Ekerot, 2006; van Beugen et al., 2013). Given the large number of PFs impinging on to a PuC, the fluctuations in basal rate are as big as the event-related high-frequency bursts. As our analysis shows, if PF synapses were static, the PuC would not be able to discriminate between high frequency bursts and background fluctuations. However, PF synapses show short-term facilitation when targeting PuC and short-term depression when targeting Basket cells (Atluri and Regehr, 1996; Bao et al., 2010; Blackman et al., 2013). Basket cells perform strong, phasic somatic inhibition to PuCs (Jörntell et al., 2010). This circuit motif closely matches with the FFI circuit investigated in the Figure 7. Based on these similarities, we argue that one of the functional implications of the specific properties of STP is to enable the PuC to discriminate between information encoded in high frequency bursts and background fluctuations.

In the neocortex, the population code in the layer 2/3 of the somatosensory (De Kock and Sakmann, 2008) and visual cortex of rats (Greenberg et al., 2008) and mice (Rochefort et al., 2009) is believed to be sparse (Petersen and Crochet, 2013), with short-lived bursts (usually < 20*ms*) of high firing rates occurring over low rate spontaneous activity (< 0.5*Hz*). Additionally, it has been recently found that pyramidal cells at layer 2/3 of the mouse somatosensory cortex show short-term facilitation when targeting cells at layers 2/3 and 5 (Lefort and Petersen, 2017). These characteristics suggest that the mechanism to discriminate between weak signals and background fluctuations may also be present in the neocortex. It is believed that such sparse representation at superficial cortical layers indicates strong stimulus selectivity (Petersen and Crochet, 2013), in which case the transient gain, provided by the target-dependent STP configuration of local pyramidal neurons, would be a suitable property for inter-layer communication.

In the hippocampus, the Schaffer collaterals bringing signals from CA3 to CA1 operate under low basal firing rates with evoked bursts of high frequency activity during short periods of time (Schultz and Rolls, 1999). The synapses from pyramidal cells in CA3 to pyramidal cells in CA1 are facilitatory and provide this pathway with extra gain control (Klyachko and Stevens, 2006). Simultaneously, Schaffer collaterals synapses to CA1 stratum radiatum interneurons show larger release probability than to pyramidal neurons (Sun et al., 2005). It is, therefore, likely that this STP-based stimulus/noise discrimination mechanism is also used to improve, transmission of sequential activity from CA3 to CA1.

As we have pointed above, STP configuration in the neocortex, hippocampus and cerebellum are consistent with the configuration that enables the neural networks to take advantage of sparse coding. It is however, not a global feature of STP in the brain, and it is important to notice that facilitating excitatory inputs to other inhibitory cells also exist in the aforementioned circuits. These facilitatory inputs mostly target interneurons that form synapses on distal dendrites (see the next paragraph). The presence of facilitatory excitatory drive to these classes of inhibitory neurons is, however, unlikely to counteract the distribution-dependent transient gains, because they produce weaker, slower and persistent dendritic inhibition. Consistent with this idea, only parvalbumin-expressing neurons (that synapse on the soma), but not somatostatin-expressing neurons (that synapse on distal dendrites), modulate stimulus response gain (Wilson et al.,2012).

The initial release probability is the most distinguishable STP parameter between Schaffer collaterals synapses onto CA1 pyramidal cells versus CA1 interneurons (Sun et al., 2005). In line with that, our approach predicts that facilitatory mechanisms that steadily increase a low initial release probability during a fast sequence of spikes (low *U*) will have a greater impact on the optimal *OD* and gain amplitude than mechanisms for fast replenishment of resources (low *τ_rec_*). However, the speed of recovery has been shown to be itself an activity-dependent feature (Fuhrmann et al., 2004;Crowley et al., 2007; Valera et al., 2012; Doussau et al., 2017) and this could in principle increase the relevance of *τ_rec_*.

The facilitatory or depressing nature of STP depends on the postsynaptic neuron type (Reyes et al., 1998; Markram et al., 1998; Rozov et al., 2001; Sun et al., 2005; Pelkey and McBain, 2007; Bao et al., 2010; Blackman et al., 2013;Larsen and Sjöström, 2015;Éltes et al., 2017). Target-dependent STP is a strong indication that such short living dynamics are relevant for specific types of information processing in the brain (Naud and Sprekeler, 2018). Here we predict that, when accompanied by specific arrangements of target-dependent STP found experimentally in different brain regions, disynaptic inhibition could further increase the gain of sparse over dense distributions and make it robust even at higher basal activity, when most of the gain possible at facilitatory excitation vanishes.

Disynaptic inhibition following excitation is a common motif throughout the brain, and different classes of inhibitory neurons are believed to serve distinct computations within their local circuits (Wilson et al., 2012; Jiang et al., 2015). Despite a wide diversity of inhibitory cell types, a classification of FF-I into two main types, perisomatic- and dendritic-targeting, seems to be coherent with findings throughout the central nervous system. A remarkable attribute of this configuration is the consistency of the short-term dynamics of excitatory synapses across local circuits: depressing to perisomatic- and facilitating to dendritic interneurons (Sun et al., 2005; Bao et al., 2010; Blackman et al., 2013; Élteset al., 2017).

Disynaptic inhibition has been implicated in controlling the precision of a postsynaptic neuron’s response to brief stimulation in the cerebellum (Mittmann et al., 2005; Ito, 2014) and hippocampus (Pouille and Scanziani, 2001). Additionally, the combination of disynaptic inhibition with target-dependent STP has been recently associated with the ability of networks to decode multiplexed neural signals in the cortex (Naud and Sprekeler, 2018). In line with these, our results show a bimodal profile of the readout neuron response to sparse or dense input code. We also demonstrate that, coexisting with the sustained gain during sparse code transmission, in a dense coding scenario, the system produces shorter periods (~ 10*ms*) of increased (decreased) spike probability right after stimulus onset (offset) (Figure 6B, gray line). This results from inhibitory conductances (GABA) which are slower than the excitatory conductances (AMPA). This very short period of firing rate modulation might work as an indication of a widespread baseline rate change in the presynaptic population.

We showed how the activity distribution of an input can exploit the nonlinearities of short-term synaptic plasticity and, with that, the theoretical potential of synaptic dynamics to endow a postsynaptic target with activity discrimination capabilities. Such mechanisms have the advantage of being in-built in synapses, not requiring further recurrent computation or any sort of supervised learning to take place. This feature is likely to be present in different brain regions, from the cerebellum to the hippocampus, and might have critical implications for general information processing in the brain.

Experimentally, these results can be tested by measuring the distribution of evoked firing rates of the neurons and STP properties of the synapses in the same brain area. Recent technological advances in stimulation systems, allowing for sub-millisecond manipulation of single and multiple cells spike activity, might soon provide means for fine control of population spike codes in intact tissues (Shemesh et al., 2017). These, together with refined methods for single cell resolution imaging of entire populations (Xu et al., 2017; Weisenburger and Vaziri, 2018) might soon allow for scrutinizing the extent of which the proposed synaptic mechanisms for distribution-dependent gain are present in neural networks.

## Materials and Methods

### Model of short-term plasticity

One parsimonious and yet powerful mathematical description of short-term synaptic dynamics was proposed already 20 years ago (Tsodyks and Markram, 1997). The Tsodyks-Markram (TM) model could first account for activity-dependent synaptic depression observed in pairs of neocortical pyramidal neurons and was soon extended to cover for facilitation (increase in probability) of vesicle release (Tsodyks et al., 1998). With a small set of parameters, the TM model is able to explain the opposed effects of depletion of available synaptic vesicles and of the increase in release probability caused by accumulation of residual calcium in the presynaptic terminal, making it suitable as a framework to conjecture general impact of STP in neural information processing.

Here, we use the TM model (equation 4) to describe the short-term synaptic dynamics. The effect of STD is modeled by depletion of the proportion of available resources, represented by the variable *x* (0 ≤ *x* ≤ 1), which instantaneously decreases after each spike and returns to 1 with recovery time *τ_rec_*. The gain effect of short-term facilitation is modeled by the facilitation factor *U* (0 ≤ *U* ≤ 1), which accounts for the accumulation of calcium at the presynaptic terminal after the arrival of an action potential. *U* transiently increases the release probability *u* (0 ≤ *u* ≤ 1), which returns to 0 with time constant *τ_f_*:

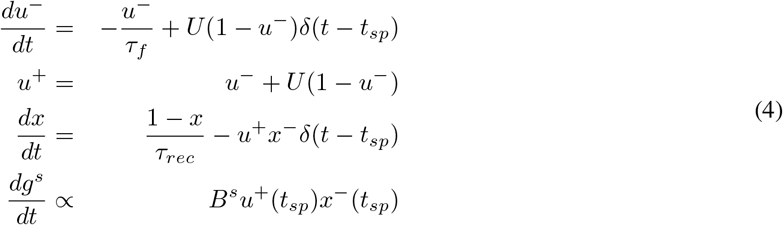

where *t_sp_* is the last spike time.

### Proportion of released resources (*PRR*)

The change in the postsynaptic conductance *g^s^* after a presynaptic spike is proportional to the instantaneous proportion of released resources (*PRR*(*t_sp_*) ∝ *u*^+^(*t_sp_*)*x*^−^(*t_sp_*)) and to the absolute synaptic strength *B^s^*. The average instantaneous PRR of a presynaptic unit can also be described as a function of a time-dependent Poissonian firing rate *r*(*t*) (Tsodyks et al., 1998) as:

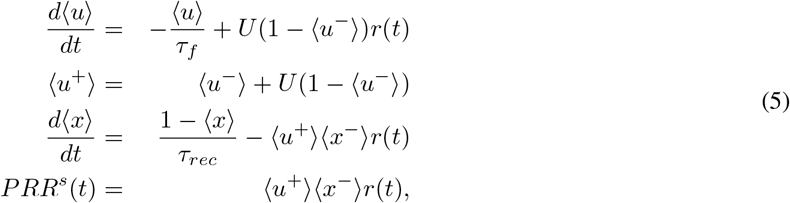

where the brackets denote the average over many realizations.

### Total effective input to a postsynaptic neuron

The total *PRR* contribution of a single synapse for a time window of duration *T_s_* is then obtained by integrating equation 5 over this period:

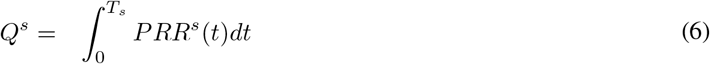

In the event of arrival of stimulus encoded as extra firing rate *R_ext_* in a homogeneous presynaptic population of size *N*, a number *N_ext_* of units will carry the stimulus encoded as a firing rate increase by *r_ext_* = *R_ext_*/*N_ext_* while the remaining units will keep their basal activity *r_bas_*. The average proportion of resources released to a target neuron by the entire population, during *T_s_*, will then be

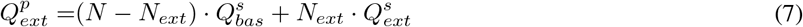

where 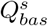 and 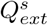 are the total *PRR* delivered by a stationary unit (firing at *r_bas_*) and a stimulus encoding unit (firing at *r_bas_* + *r_ext_*), respectively.

### Gain in the effective input

We quantify the gain in 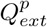 for a given *N_ext_* relative to the 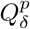 caused by an input of the same intensity but with dense distribution (when *N_ext_* = *N*) as

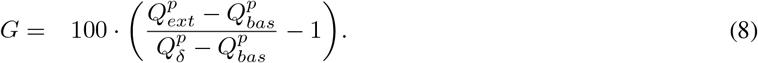

*G* is a non-monotonic function of *N_ext_* with a single maximum value *G_max_* at *N_opt_* (see Figure 2D).

### Optimal distribution

The optimal distribution of the activity (*OD*) is defined as the fraction of the optimal number of encoding units *N_opt_* in a given population of size *N*, that is, *OD* = *N_opt_*/*N*. Because the optimal code, *N_opt_* = *R_ext_*/*r_opt_*, is the distribution that maximizes the gain over the dense distribution with the same input magnitude, *N* = *R_ext_*/*r_δ_, OD* can be written as

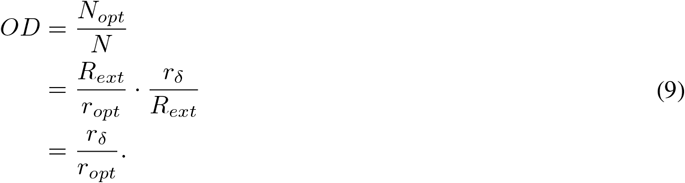

Because *r_δ_* is defined as a fraction of *r_bas_* and *r_opt_* is fixed given the STP parameters and *r_bas_*, an interesting consequence is that *OD* becomes independent of any particular choices of *N* and *N_opt_*. Since the optimal encoding rate is constrained by *r_δ_* < *r_opt_* < ∞, the optimal distribution will be constrained to 0 < *OD* < 1 (see Figure 3D), with values close to zero or one characterizing, respectively, sparse or dense distributions.

### Optimum rate (*r_opt_*) and maximum gain (*G_max_*) estimation

Equation 8 describes the gain *G* obtained by encoding a stimulus *R_ext_* into *N_ext_* units (with rates increased by *r_ext_* = *R_ext_*/*N_ext_*) as opposed to *N* units (with rates increased by *r_δ_* = *R_ext_*/*N*). The peak of this function (*G_max_*) is achieved by an optimum number of encoding units *N_opt_* with their rate increased by *r_opt_* = *R_ext_*/*N_opt_*. This maximum point can be found by taking the derivative of the gain function with respect to *r_ext_* and setting it equal to zero,

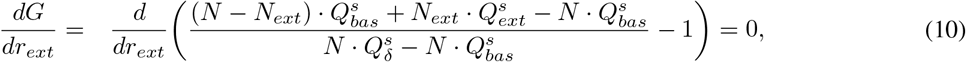

this can be further simplified into

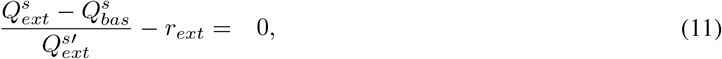

where 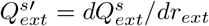 and *r_ext_* that solves the equation is denominated *r_opt_*. This solution is independent of the stimulus intensity *R_ext_* and population size *N*, resulting in the iso-frequency line in Figure 2F.

For the optimum rate *r_opt_*, the gain (equation 8) can be written as

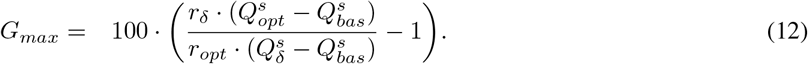

Assuming that 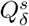 is linear with slope *S^s^* for small *r_δ_*, that is, 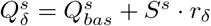 (See *Linear approximation of* 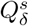, below), then *G_max_* can be further simplified into

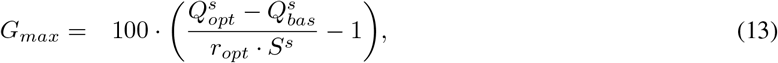

which makes *G_max_* independent of the stimulus intensity *R_ext_* and population size *N*.

### Combined optimum rate 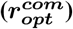 and maximum gain 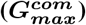 estimation

When an axon branches to connect to different targets, STP properties might be target dependent. In the case of excitatory fibers driving FF-EI motifs, with *s*1 directly exciting a readout neuron and *s*2 driving the local FF-I circuit, the gain is

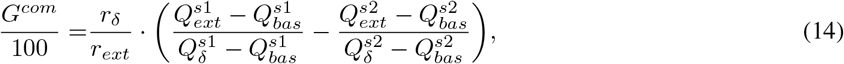

Taking the derivative of *G^com^* with respect to *r_ext_*, setting it equal to zero and assuming again that 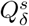 is linear with slope *S^s^* for both synapses, it yields

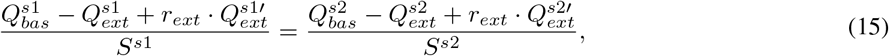

for which the solution, 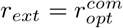, is independent of the stimulus intensity *R_ext_* and population size *N*. The optimum combined gain is then

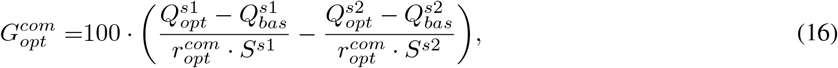

which is also independent of the stimulus intensity *R_ext_* and population size *N*.

### Linear approximation of 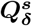

We solve *Q^s^* numerically (equation 6) and show that it behaves linearly for a moderate range of rates in different STP regimes (Figure 8). The approximation by a linear function, 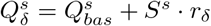, allows *G_max_* to be independent of the stimulus intensity and population size (equation 13).

**Figure 8:**
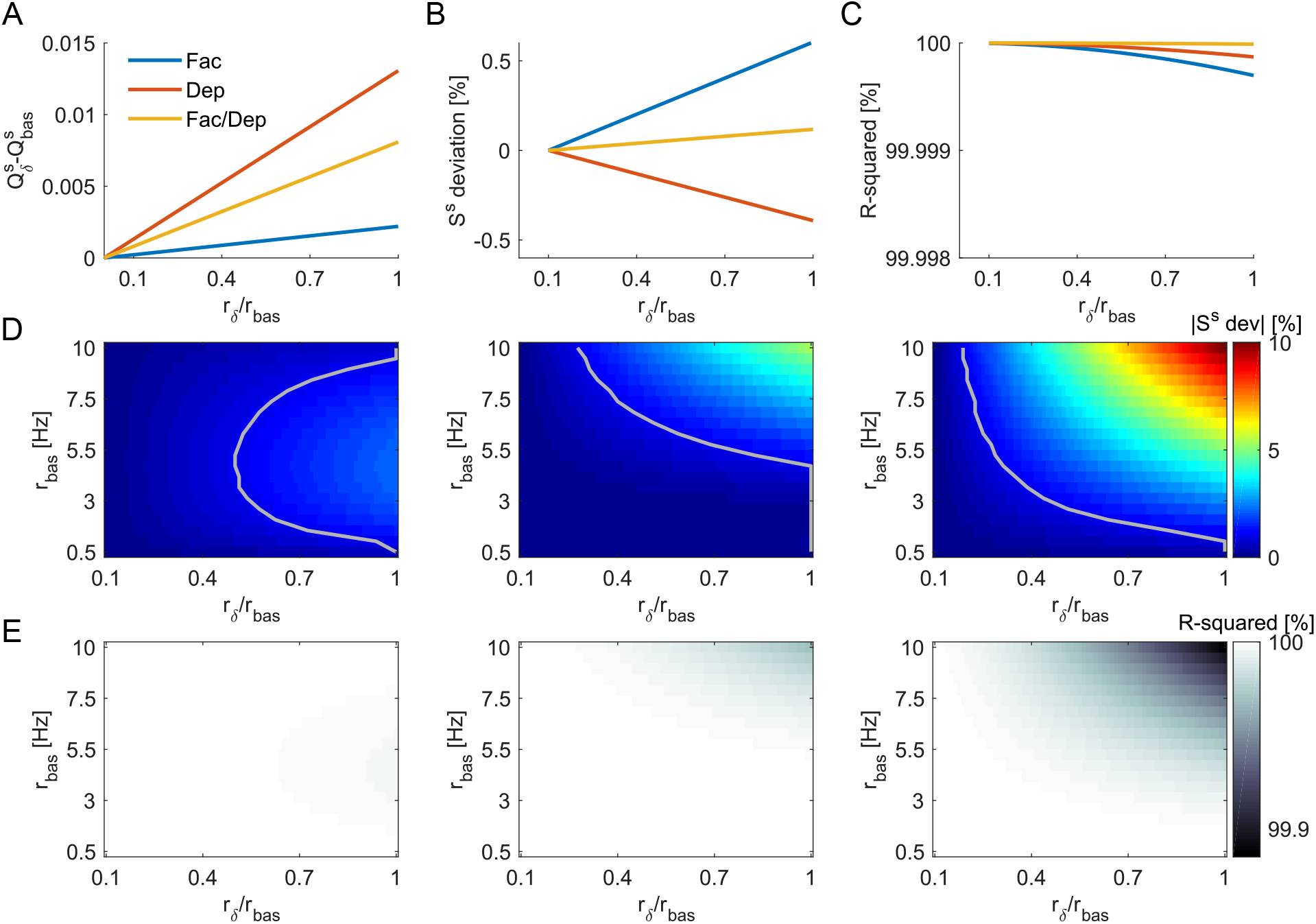
Linear approximation of 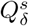. **A** For basal rate *r_bas_* = 0.5*Hz* and *T_s_* = 40*ms*, 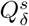 is approximately linear for all STP regimes, facilitatory (*U* = .1,*τ_rec_* = 50*ms*,*τ_f_* = 2000*ms*), facilitatory/depressing (*U* = .4,*τ_rec_* = 100*ms*,*τ_f_* = 1000*ms*) or depressing (*U* = .7,*τ_rec_* = 2000*ms*,*τ_f_* = 500*ms*). **B** Slope deviation for increasing *r_δ_* in comparison to the slope for *r_δ_* = 0.1 · *r_bas_* is always smaller than 0.6% for the three synapses. **C** R-squared is always close to 100% for the three synapses. **D** Absolute value of slope deviation, similar to panel B, but for *r_δ_* departing from several different values of *r_bas_*. The gray line marks |*S^s^* dev| = 1%. We observe that the linear approximation will work well throughout a large space (left from the gray line) for facilitation (*left*) and facilitation/depression (*middle*)regimes, and gets a bit more constrained for depressing (*right*) synapses. **E** Similar to panel C, but for *r_δ_* departing from several different values of *r_bas_*.

To which extent is the linear approximation valid? To investigate this, we solve 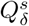 for gradually increasing *r_δ_* departing from a range of different basal levels *r_bas_* = 0.5…10*Hz*. We then compare the slopes for each *r_δ_* to the slope for *r_δ_* = 0.1 · *r_bas_* and see how much they deviate from it (Figure 8B,D). If, for a given *r_bas_*, increasing *r_δ_* would result in significant change in the regressed *S^s^*, then *G_max_* would be dependent on the stimulus intensity *R_ext_*. We also show the R-squared statistics to confirm the accuracy of the linear approximation (Figure 8C,E).

As we observe, for low signal-to-basal ratios (*r_δ_*/*r_bas_* < 1), there is a wide range of rates for which the approximation is good enough, with |*S^s^* dev| < 1% and R-squared> 99.9%. Specially for low *r_bas_*, the approximation is valid for the whole length of *r_δ_*.

### Readout neuron model

We make our proof of concept with a simulated conductance-based integrate-and-fire (IF) neuron with its membrane voltage *V_m_* described by

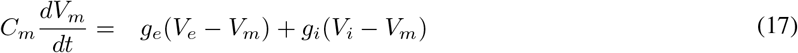

where *C_m_* = 250*pF* is the membrane capacitance, *g_e_* and *g_i_* are respectively the excitatory and inhibitory input conductances and *V_e_* = 0*mV* and *V_i_* = −75*mV* are the excitatory and inhibitory synaptic reverse potentials. When a spike occurs, the membrane voltage is reset at *V_reset_* = −60*mV* and held at this value for a refractory period of 2*ms*. The synapses were modeled by *α*-functions (Kuhn et al., 2004) with time constants *τ_e_* = .5*ms* for excitatory and *τ_i_* = 2*ms* for inhibitory synapses.

### Feedforward inhibitory circuit and input

The presynaptic population consisted of *N* = 160000 units that connected to the postsynaptic neuron in a feedforward excitation-inhibition arrangement (Figure 1C). The population stationary basal rate was *R_bas_* = 80*kHz*, with the individual basal rate *r_bas_* = 0.5*Hz*.

At the stationary basal rate, the synaptic states are described by

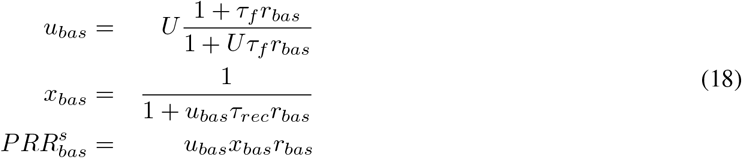

where 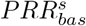 is the expected rate of proportion of released resources by each synapse with STP parameters {*U, τ_rec_, τ_f_*}.

We simulate a neuron that, during stationary basal activity, is kept in the fluctuation-driven regime through excitation-inhibition input balance (Kuhn et al., 2004). While excitation is provided directly by *s*1, disynaptic inhibition is modulated by *s*2 in a linear fashion,

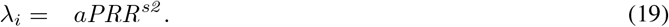

The inhibitory firing rate that keeps the target neuron membrane potential fluctuating around the mean value of *μ*(*V_m_*) during stationary basal activity can be approximated by a linear function of the excitation (adapted from Kuhn et al.(2004)),

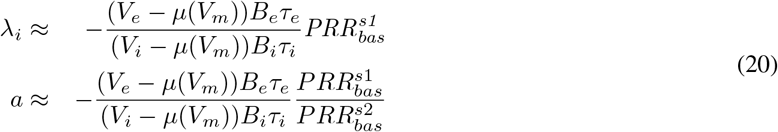

where *B_e_* and *B_i_* are the maximum amplitudes for the excitatory and inhibitory synaptic conductances. Equation 20 allows to find the linear scale of equation 19 that fulfill the condition *μ*(*V_m_*) = −53*mV*. The inhibitory synapses are kept static (no STP). The extra presynaptic activity happens in blocks of *T_s_* = 40*ms* and is defined as *sparse* (*N_ext_* = *N_opt_*) or *dense* (*N_ext_* = *N*).

All the analyses, simulations and figures were performed in Matlab and Python. The IF model simulations were performed using Euler’s method with time step of 0.1*ms* implemented in the neural simulator Brian2 (Stimberg et al., 2014).

### Limitations and possible extensions

The transient enhancement or depression of synaptic efficacy by presynaptic mechanisms consists of many independent processes (Zucker and Regehr, 2002). The TM model is a tractable and intuitive way to account for these two phenomena of interest, but this parsimony comes at the cost of biophysical simplifications. For example, it assumes the space of available resources is a continuum (0 < *x* < 1) as opposed to the known discrete nature of transmitter-carrying vesicles. However, we argue that when modeling a large number of simultaneously active synapses, the variable of interest (population *PRR*) can be approximated by a continuous variable. The nonuniform amount of transmitters per vesicle might further contribute to the validity of this assumption.

Detailed STP models that try to account for specific intracellular mechanisms were already proposed in the literature (Dittman et al., 2000), also accounting for the stochasticity of the release process (Sun et al., 2005; Kandaswamy et al.,2010). We argue that, although the quantitative results could possibly differ with more complex models, the qualitative outcome of our analysis would remain: that presynaptic short-term facilitation (depression) yields a substantial positive (negative) gain to sparse over dense population codes. Nevertheless, it would be interesting to see how the gain and optimum rate predictions would be shaped by more detailed models.

Our analyses do not account for use-dependent recovery time, changes in the readily releasable pool size (Kaeser and Regehr, 2017) or vesicles properties heterogeneity. The effects of postsynaptic receptor desensitization and neurotransmitter release inhibition by retrograde messengers (Brown et al., 2003) are likely to decrease the estimated gains by counteracting facilitation. Another interesting extension could be done with multi-compartment neuron models, which could be used to further investigate the effects of input STP heterogeneity at compartment-dependent input (Vetter et al., 2001; Grillo et al., 2018).

If the same patterns of bursts tend to happen repeatedly (e.g. parallel fibers in cerebellum during continuously repetitive movement), there might be an optimum inter-burst interval *IBI^opt^* for which, if bursts happen faster than *IBI^opt^*, the signal would be compromised (by a not fast enough vesicles recovery time) and if bursts happen separated by intervals longer then *IBI^opt^* no extra gain will happen. Experimental evidence points to resonance at the theta-frequency oscillations range (5.0 ~ 10.0*Hz*, 100 ~ 200*ms* IBI) as important for cortical-cerebellar drive (Gandolfi et al., 2013;Chen et al., 2016) and hippocampus (Buzsáki, 2002). In these cases, the slower interaction between different pools of vesicles (Rizzoli and Betz, 2005) are likely to play a role in information transfer. Augmentation, a form of transient synaptic enhancement that can last for seconds, is also likely to play a role in these cases (Kandaswamy et al., 2010;Deng and Klyachko, 2011).

## Acknowledgments

We would like to thank Dr. Gilad Silberberg, Dr. Erik Fransén, Dr. Philippe Isope and Martino Sindaci for helpful suggestions and feedback. This work was funded in part by: the EU Erasmus Mundus Joint Doctorate Program EUROSPIN, The International Graduate Academy (IGA) of the Freiburg Research Services (to Luiz Tauffer), Swedish Research Council (Research Project Grant, StratNeuro, India-Sweden collaboration grants to Arvind Kumar).

